# Improving cell-type identification with Gaussian noise-augmented single-cell RNA-seq contrastive learning

**DOI:** 10.1101/2022.10.06.511191

**Authors:** Ibrahim Alsaggaf, Daniel Buchan, Cen Wan

## Abstract

Cell-type identification is an important task for single-cell RNA-seq (scRNA-seq) data analysis. In this work, we proposed a novel Gaussian noise augmented scRNA-seq contrastive learning framework (GsRCL) to learn a type of discriminative feature representations for cell-type prediction tasks. The experimental results suggest that the feature representations learned by GsRCL successfully improved the accuracy of cell-type prediction using scRNA-seq expression profiles.

## Introduction

Single-cell sequencing methods aim to characterise the DNA or RNA contents of single cells using current Next Generation Sequencing technologies. Typically, the experimental target is to seek to characterise the DNA or transcriptome of target cells^1^. In transcriptome analysis, expression profiles of the cell’s genes can be measured by amplifying a cell’s RNA transcripts, and then sequencing the population of nucleotide sequences – usually referred to as single-cell RNA sequencing methods or scRNA-seq. Such experiments yield information about which genes are being transcribed and RNA concentrations can give proxy estimates of the amount of the protein being produced for each transcribed gene^2^. The expression profiles provide crucial information for understanding complex biological problems, such as gene-regulatory logic^3,4^, genomic function^5,6^, and cell-type identification^7,8^. Cell-type identification rests on the observation that differing cell-types have specific gene expression profiles. Therefore, expression profiles can be used to identify cell-types in tissue samples^9^, and unique expression profiles have even been used to uncover previously unknown cell-types within tissues^10^. Due to the importance and utility of cell-type identification a number of automated methods have already been produced such as scPred^11^, scAnnotatR^12^, scTYPE^13^ and scSimClassify^14^. In the following paper we discuss the use of contrastive learning in this task.

Contrastive learning has recently drawn a great attention given its success in achieving state-of-the-art performance over a variety of predictive tasks, e.g. image classification^15,16^ and speech recognition^17,18^. In general, contrastive learning is a type of self-supervised learning paradigm where the task is to learn a model that encodes the similarity information between instances without considering their labels. In the model’s latent space, the instances that are similar to each other are projected to a proximal region, whereas those differing instances are projected into areas that are far away from each other. In order to learn this type of model, the classic frameworks like SimCLR^15^ and SupCon^19^ adopt data augmentations to create different views of the original training instances. These views are used to learn neural network models that aim to maximize the similarity between views that belong to the same original instance. Therefore, data augmentation is a key component of the contrastive learning paradigm. Recent works in computer vision contrastive learning^15,19–22^ adopt spatial transformations (i.e. random cropping, random flipping, and rotation), and colour transformations (i.e. colour distortion, Gaussian blur, and gray scale) to create views of target images. When handling text data, paraphrasing and word replacement are adopted as data augmentation methods^23^. Other data augmentation methods like node dropping, edge perturbation, graph subsampling, and subgraph removal are also proposed for graph-specific contrastive learning tasks^24,25^. Note that most of the existing data augmentation methods exploit structured information retained in the data, but cannot be applied on data that do not bear such structured information, e.g. scRNA-seq data.

Contrastive learning has already been applied to a variety of unsupervised learning-based scRNA-seq data analysis tasks, e.g. scRNA-seq data clustering, scRNA-seq data batch effects removal, and scRNA-seq data integration. SMILE^26^ previously proposed to use Gaussian noise to create augmentations of the biological data, which are integrated and projected into a latent space so similar cells can be clustered by using an appropriate contrastive learning loss function. GLOBE^27^ also propose a contrastive learning-based scRNA-seq integration method that exploits nearest neighbour cells’ expression profiles to create different data views. HDMC^28^ have proposed a batch effects removal method that exploits contrastive learning loss to work with pre-defined similar cell pairs and noisy cell pairs without changing the distribution of the original gene expression profiles. Contrastive-sc^29^ and scNAME^30^ both adopt a random masking strategy, i.e. masking an arbitrary set of genes where they are ignored from the underlying computation, to create different views of target gene expression profiles to work with contrastive learning loss function in order to cluster scRNA-seq data. Lastly we note, CLEAR^31^ use a combination of data augmentation methods, i.e. random masking, Gaussian noise replacement, random swapping, and crossover to create different views for target cells in order to integrate scRNA-seq data using contrastive learning approaches.

In this work, we proposed a novel cell-type identification method, namely GsRCL, which exploits the well-known Gaussian noise to augment the original gene expression profiles in order to learn contrastive learning feature representations. Analogous to the work^26^, we also use Gaussian noise to create views for different target cells’ expression profiles but then train a feature extractor to transform the gene expression profiles for the downstream supervised learning tasks, i.e. cell-type identification. As almost all scRNA-seq contrastive learning methods proposed so far focus on unsupervised learning tasks, it is timely to investigate the performance of this Gaussian noise data augmentation method on contrastive learning and downstream supervised learning tasks, due to its simplicity and the concern on the well-known overfitting issue – the effect of the contrastive learning-based feature extractor against the downstream classifier is still unknown. The experimental results suggest that our GsRCL method successfully improved the accuracy of cell-type identification and outperformed the random genes masking data augmentation method^29^. A further analysis reveals that the self-supervised contrastive learning loss has higher robustness than the supervised contrastive learning loss against the overfitting issue.

## Results

### Overview of GsRCL

In general, the proposed GsRCL method consists of two stages of training. As shown in Figure 1.a, the first stage of training aims to learn a feature extractor that generates a type of discriminative feature representations using a Gaussian noise augmented contrastive learning approach. Given one gene’s *d*-dimensional scRNA-seq expression profiles *s*, GsRCL firstly creates two *d*-dimensional vectors *v*_1_ and *v*_2_, where the value for each dimension is randomly drawn from the Gaussian distribution 𝒩 (0, 1). Then those two different Gaussian noise vectors are respectively added with the *d*-dimensional scRNA-seq expression profiles in order to create two views 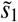 and 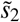 for the contrastive learning task, i.e. 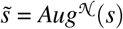 = *Aug*^𝒩^ (*s*). Those two views are then used as inputs for an encoder 𝒢 (a.k.a. the feature extractor), which generates feature representations inputs for a projector head ℋ. The projector head maps the views into a latent space, i.e. *z* = ℋ (𝒢 (*Aug*^𝒩^ (*s*))), where *z* denotes the latent space feature representations of one gene’s original scRNA-seq expression profiles. Then those two views are pulled closer to each other and pushed further away from all other genes in the given dataset by using the contrastive learning loss (see Methods).

**Figure 1.**
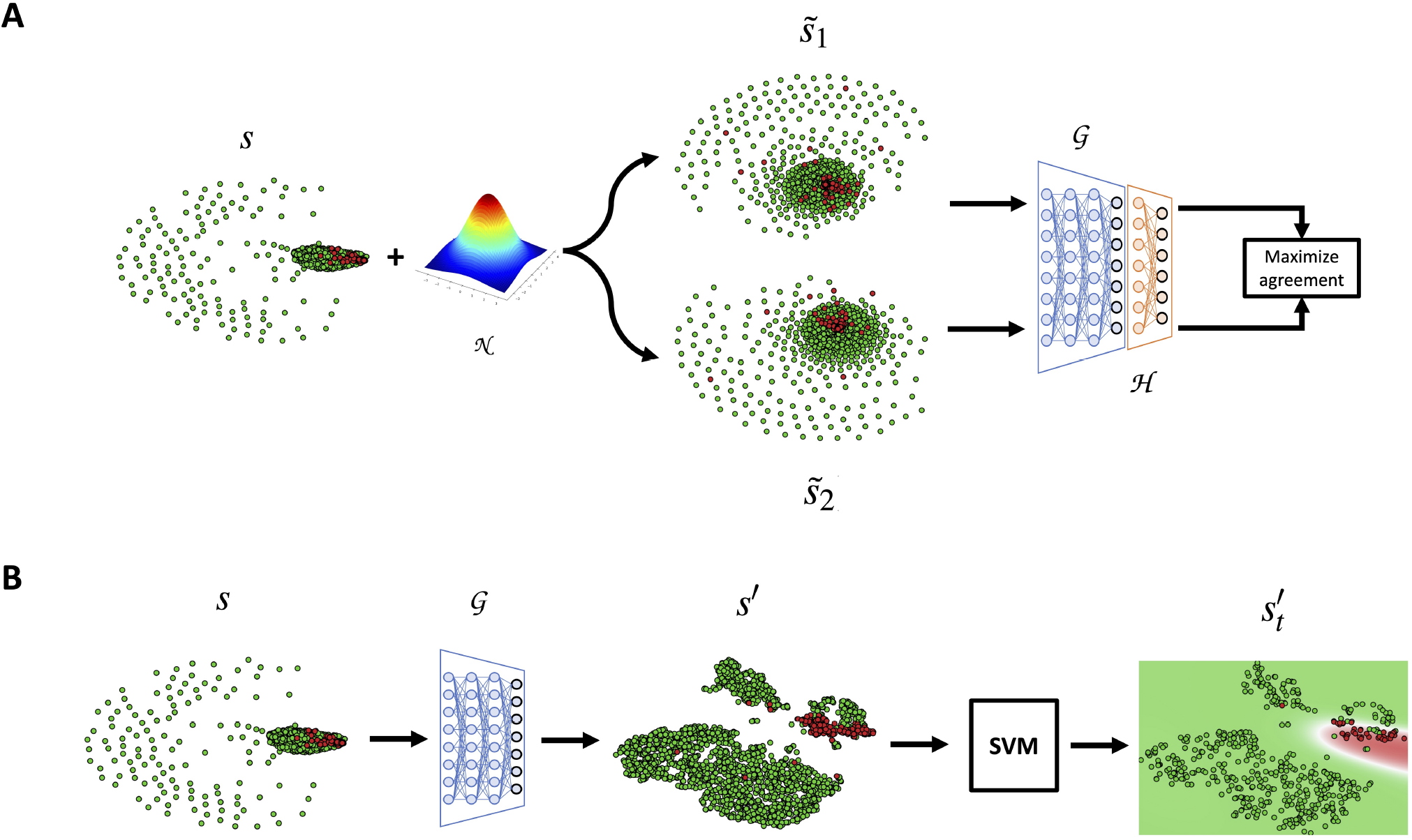
The flowchart for GsRCL. The GsRCL method consists of two stages of training. (a) The first stage is to use Gaussian noise 𝒩 to create two views 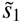 and 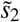 of the original input scRNA-seq expression profiles *s*. These two new views are encoded by an encoder 𝒢 and then projected into a latent space by a projector head ℋ. Those two projected feature representations are pushed closer in the latent space by the contrastive learning loss. (b) After finishing the stage (a) training, the encoder 𝒢 is used to generate contrastive learning feature representations *s*^′^ of the original input scRNA-seq expression profiles *s*. The generated feature representations *s*^′^ are then used to train a support vector machine (SVM) to predict the encoder 𝒢 transformed cell-types of unlabeled scRNA-seq expression profiles 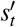.

The second stage of training aims to use the generated discriminative feature representations to predict cell-types. During the first stage of contrastive learning, we use an SVM classifier and a validation dataset to select the optimal encoder whose generated feature representations lead to the highest predictive accuracy. Then the selected optimal encoder is used to transform the original scRNA-seq expression profiles in the training and testing datasets. As shown in Figure 1.b, the original scRNA-seq expression profiles are successfully transformed as a new type of feature representations where the genes bearing two different class labels are grouped separately, leading to an improved predictive accuracy, since a better decision boundary can be learned. Note that, any testing genes’ scRNA-seq expression profiles also need to be transformed by using the trained encoder before being used as inputs for the SVM, which was trained by using the transformed training genes’ scRNA-seq expression profiles.

We evaluated the performance of the proposed GsRCL method by using self-supervised^26^ and supervised contrastive learning^19^ losses, respectively. Table 1 display the Matthews correlation coefficient and average precision values obtained by feature representations learned by different data augmentation methods working with self-supervised and supervised contrastive learning losses over 13 peripheral blood mononuclear cells (PBMC) datasets. The well-known support vector machine was adopted as the classifier.

**Table 1.** The performance in identifying cell-types using different contrastive learning feature representations and the original scRNA-seq expression profiles.

| Protocol of<br>PBMC datasets<br>(Positive class) |  | Self-supervised contrastive learning loss |  |  |  |  | Supervised contrastive learning loss |  |  |  |  | Benchmark<br>(Original<br>expression<br>profiles) |
| --- | --- | --- | --- | --- | --- | --- | --- | --- | --- | --- | --- | --- |
|  |  | Gaussian<br>Noise | Random<br>masking<br>500 | Random<br>masking<br>1000 | Random<br>masking<br>3000 | Random<br>masking<br>5000 | Gaussian<br>Noise | Random<br>masking<br>500 | Random<br>masking<br>1000 | Random<br>masking<br>3000 | Random<br>masking<br>5000 |  |
| 10Xv2<br>(Natural killer cell) | MCC | **<u>0.797</u>** | 0.167 | 0.170 | 0.662 | 0.757 | **0.788** | 0.138 | 0.191 | 0.641 | 0.699 | 0.778 |
|  | AP | **<u>0.908</u>** | 0.208 | 0.267 | 0.839 | 0.874 | 0.839 | 0.141 | 0.250 | 0.758 | **0.841** | 0.905 |
| 10Xv2<br>(Cytotoxic T cell) | MCC | **<u>0.887</u>** | 0.426 | 0.586 | 0.817 | 0.825 | **0.871** | 0.416 | 0.556 | 0.779 | 0.798 | 0.855 |
|  | AP | **<u>0.970</u>** | 0.643 | 0.760 | 0.941 | 0.942 | **0.942** | 0.633 | 0.742 | 0.849 | 0.861 | 0.958 |
| 10Xv2<br>(CD4<sup>+</sup> T cell) | MCC | **<u>0.905</u>** | 0.373 | 0.614 | 0.863 | 0.862 | **0.882** | 0.344 | 0.559 | 0.841 | 0.857 | 0.879 |
|  | AP | **<u>0.972</u>** | 0.572 | 0.784 | 0.951 | 0.952 | 0.937 | 0.536 | 0.718 | 0.934 | **0.949** | 0.960 |
| 10Xv3<br>(Natural killer cell) | MCC | **<u>0.855</u>** | 0.295 | 0.527 | 0.839 | 0.796 | **0.836** | 0.342 | 0.554 | 0.798 | 0.793 | 0.844 |
|  | AP | 0.922 | 0.345 | 0.672 | **0.948** | 0.929 | 0.937 | 0.372 | 0.659 | **0.949** | 0.932 | 0.938 |
| 10Xv3<br>(Cytotoxic T cell) | MCC | 0.879 | 0.482 | 0.686 | **<u>0.893</u>** | 0.878 | **0.888** | 0.433 | 0.646 | 0.877 | 0.858 | 0.871 |
|  | AP | 0.976 | 0.657 | 0.812 | **<u>0.982</u>** | 0.980 | **0.977** | 0.631 | 0.796 | 0.961 | 0.915 | 0.974 |
| Drop-Seq<br>(CD4<sup>+</sup> T cell) | MCC | **<u>0.919</u>** | 0.578 | 0.620 | 0.826 | 0.867 | **0.898** | 0.544 | 0.569 | 0.816 | 0.843 | 0.896 |
|  | AP | **<u>0.985</u>** | 0.749 | 0.771 | 0.957 | 0.964 | **0.961** | 0.705 | 0.725 | 0.937 | 0.960 | 0.981 |
| Drop-Seq<br>(Cytotoxic T cell) | MCC | **<u>0.867</u>** | 0.483 | 0.549 | 0.774 | 0.789 | **0.864** | 0.458 | 0.475 | 0.703 | 0.740 | 0.822 |
|  | AP | **<u>0.977</u>** | 0.705 | 0.712 | 0.917 | 0.944 | **0.962** | 0.674 | 0.683 | 0.845 | 0.882 | 0.955 |
| Drop-Seq<br>(CD14<sup>+</sup> monocyte) | MCC | **<u>0.884</u>** | 0.746 | 0.776 | 0.807 | 0.760 | **0.869** | 0.636 | 0.596 | 0.530 | 0.525 | 0.828 |
|  | AP | **<u>0.961</u>** | 0.783 | 0.836 | 0.901 | 0.886 | **0.932** | 0.691 | 0.614 | 0.619 | 0.634 | 0.925 |
| inDrop<br>(CD4<sup>+</sup> T cell) | MCC | **<u>0.752</u>** | 0.550 | 0.544 | 0.684 | 0.672 | **0.719** | 0.476 | 0.465 | 0.584 | 0.578 | <u>0.762</u> |
|  | AP | **0.859** | 0.662 | 0.668 | 0.827 | 0.821 | **0.806** | 0.564 | 0.593 | 0.773 | 0.732 | <u>0.862</u> |
| inDrop<br>(CD14<sup>+</sup> monocyte) | MCC | **0.910** | 0.861 | 0.850 | 0.836 | 0.850 | **0.913** | 0.853 | 0.871 | 0.867 | 0.842 | 0.897 |
|  | AP | **<u>0.987</u>** | 0.965 | 0.961 | 0.957 | 0.956 | 0.962 | 0.958 | 0.966 | **0.968** | 0.956 | 0.986 |
| inDrop<br>(Cytotoxic T cell) | MCC | **0.819** | 0.754 | 0.761 | 0.804 | 0.788 | **0.827** | 0.748 | 0.769 | 0.809 | 0.787 | <u>0.845</u> |
|  | AP | **0.948** | 0.913 | 0.918 | 0.917 | 0.919 | 0.926 | 0.894 | 0.913 | **0.947** | 0.941 | <u>0.958</u> |
| Seq-Well<br>(Cytotoxic T cell) | MCC | **<u>0.820</u>** | 0.628 | 0.643 | 0.775 | 0.797 | **0.803** | 0.615 | 0.638 | 0.722 | 0.763 | 0.813 |
|  | AP | **<u>0.939</u>** | 0.839 | 0.841 | 0.932 | 0.938 | 0.900 | 0.821 | 0.818 | 0.901 | **0.920** | 0.938 |
| Seq-Well<br>(CD4<sup>+</sup> T cell) | MCC | **<u>0.677</u>** | 0.340 | 0.422 | 0.611 | 0.631 | **0.643** | 0.286 | 0.388 | 0.542 | 0.557 | 0.639 |
|  | AP | **<u>0.789</u>** | 0.422 | 0.548 | 0.759 | 0.776 | 0.696 | 0.402 | 0.488 | 0.689 | **0.742** | 0.772 |

### GsRCL Performance

The contrastive learning feature representations learned by the proposed GsRCL method showed the overall best predictive performance. We note these had the best overall predictive performance and improved the predictive accuracy of the original scRNA-seq expression profiles. As shown in Table 1, GsRCL (Gaussian noise augmentation with self-supervised contrastive learning loss) obtained the overall highest MCC values (as denoted by underlines) on 9 out of 13 tested datasets, and also obtained the overall highest average precision values on 8 out of 13 datasets. The GsRCL-learned feature representations also successfully improved the predictive accuracy of the original scRNA-seq expression profiles, which merely obtained the overall best predictive performance on two datasets, i.e. inDrop CD4^+^ T cell and inDrop Cytotoxic T cell. In addition, the Gaussian noise augmentation method outperformed all random genes masking data augmentation methods. The random genes masking data augmentation methods only obtained a higher MCC value on the 10Xv3 Cytotoxic T cell dataset, when working with self-supervised contrastive learning loss. However, when working with supervised contrastive learning loss, the random genes masking data augmentation methods obtained higher average precision values on 7 out of 13 datasets.

Figures 2.a and 2.b show the precision-recall curves for different data augmentation methods working with two different contrastive learning losses on classifying the Drop-seq CD14^+^ monocyte cell-type. It is obvious that the proposed Gaussian noise augmentation-based GsRCL (denoted by the black curve) covers the largest areas (i.e. 0.961 and 0.932 of average precision values for self-supervised and supervised losses, respectively) than all other compared methods. The red curve that is used to denote the benchmark (i.e. the original scRNA-seq expression profiles) cover the second largest area, suggesting that all variants of the random genes masking data augmentation methods failed to improve the predictive performance of the original scRNA-seq expression profiles. Analogously, the bar-chart shown in Figure 2.c suggests that GsRCL obtained highest MCC value (i.e. 0.884 and 0.869 for self-supervised and supervised contrastive learning losses, respectively), compared with all variants of the random genes masking data augmentation methods and the original scRNA-seq expression profiles.

**Figure 2.**
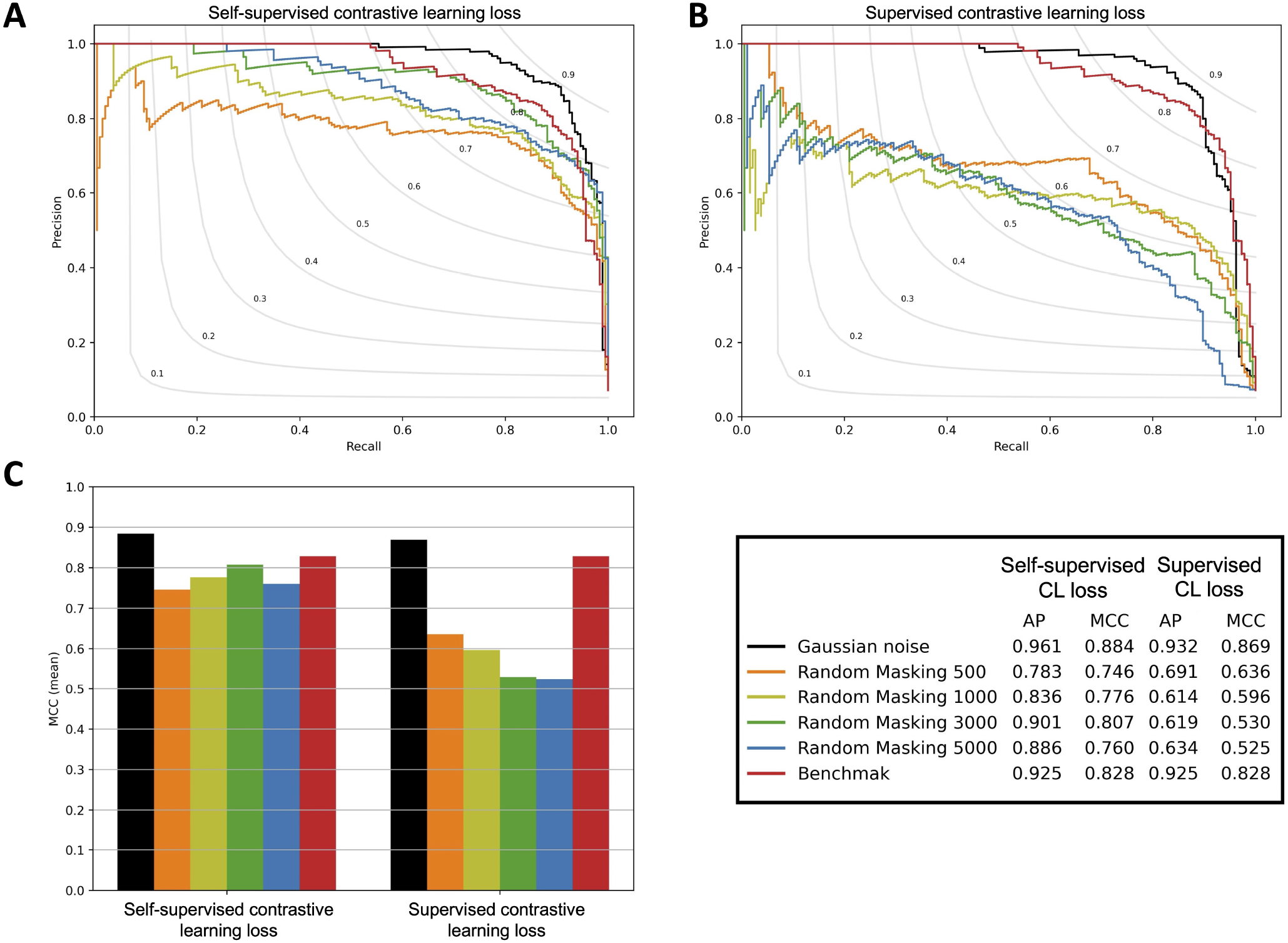
The precision-recall curves obtained by different data augmentation methods working with self-supervised and supervised contrastive learning losses for identifying the CD14^+^ monocyte cell-type.

### Self-supervised contrastive learning obtains higher accuracy and shows stronger robustness against overfitting

The feature representations learned by the self-supervised contrastive learning loss obtains higher predictive accuracy than the feature representations learned by the supervised contrastive learning loss. We evaluated different data augmentation methods for the contrastive learning tasks by using self-supervised and supervised contrastive learning losses, respectively. Table 2 shows the results of pairwise comparisons between those two losses working with each individual data augmentation method. It is obvious that all different data augmentation methods working with self-supervised contrastive learning loss obtain higher predictive accuracy on majority of datasets, suggesting that self-supervised contrastive learning loss learns better feature representations in terms of the downstream supervised learning tasks than the supervised contrastive learning loss.

**Table 2.**
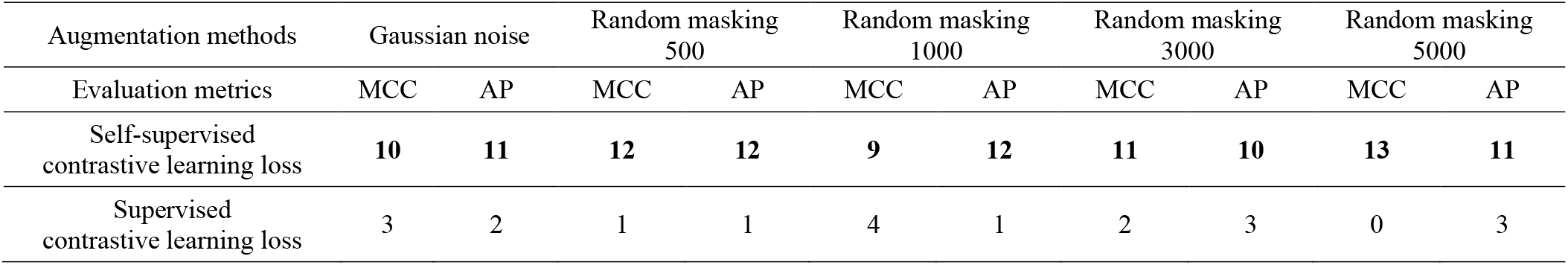
The frequency of higher predictive accuracy for each data augmentation method and two different contrastive learning loss functions.

| Augmentation methods | Gaussian noise |  | Random masking<br>500 |  | Random masking<br>1000 |  | Random masking<br>3000 |  | Random masking<br>5000 |  |
| --- | --- | --- | --- | --- | --- | --- | --- | --- | --- | --- |
|  | MCC | AP | MCC | AP | MCC | AP | MCC | AP | MCC | AP |
| Self-supervised<br>contrastive learning loss | 10 | 11 | 12 | 12 | 9 | 12 | 11 | 10 | 13 | 11 |
| Supervised<br>contrastive learning loss | 3 | 2 | 1 | 1 | 4 | 1 | 2 | 3 | 0 | 3 |

## Discussions

### Self-supervised learning shows stronger robustness against the overfitting issue

We further investigated the overfitting issue of self-supervised and supervised contrastive losses by considering the training accuracy during the down-stream SVM classifier training stage. For each fold of the cross validation, we use all training folds to train a SVM classifier, then use those training folds as inputs to obtain a training accuracy of that trained SVM classifier. The training accuracy reflects the fitting quality of the SVM classifier with regards to the training data only. We then use the testing fold to obtain a testing accuracy, which is used to evaluate the overfitting degree by considering the training accuracy. Figure 3 illustrates the distributions of the training and testing accuracy obtained by two contrastive learning losses. It is obvious that the feature representations learned by the self-supervised contrastive learning loss has stronger robustness against the overfitting issue, because it obtained lower training accuracy, but showed better predictive performance on the testing data, according to both MCC and AP values. On the contrary, the supervised contrastive learning loss obtained much higher training accuracy, but showed worse predictive performance on the testing data.

**Figure 3.**
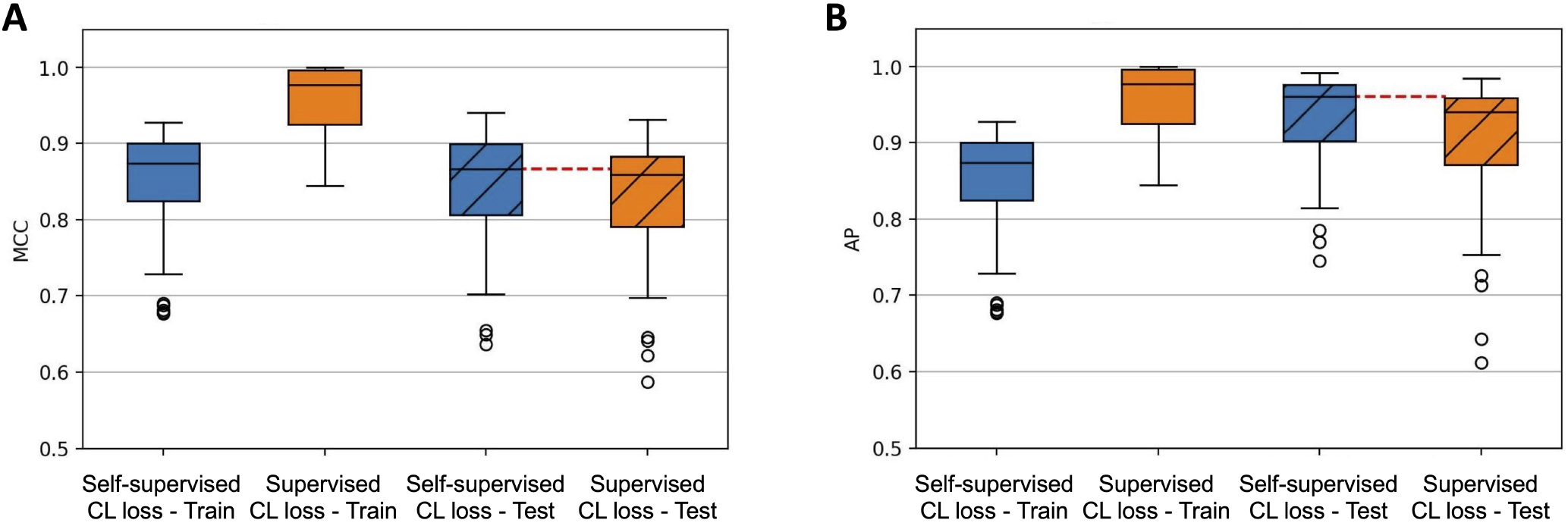
The boxplots of training and testing accuracy, i.e. MCC values (A) and AP values (B), obtained by self-supervised and supervised contrastive learning losses with Gaussian noise augmentation.

### The contrastive learning feature representations learned by GsRCL show stronger robustness against the class imbalance issue

As all those 13 PBMC datasets discussed in this work consist of class labels whose distributions are highly imbalanced, we evaluated the robustness of GsRCL (i.e. Gaussian noise augmentation working with the self-supervised loss) and the benchmark method (i.e. using the original gene expression profiles) against the class imbalance issue. We calculated the label imbalance degree *I* by Equation 1^32^, where # (Minor) denotes the number of instances belong to the minority class, and # (Major) denotes the number of instances belong to the majority class. As shown in Table 3, the label imbalance degrees of the 13 datasets range from 0.415 to 0.936. As shown in Figure 4.a, in general, the MCC values obtained by GsRCL and the benchmark method both decrease when the imbalance degree increases, but GsRCL shows stronger robustness since its MCC values are still almost all higher than the ones obtained by the benchmark method. In terms of the average precision values, both methods show a very similar pattern, as shown in Figure 4.b.

**Table 3.** The characteristics of 13 peripheral blood mononuclear cells scRNA-seq datasets used in this work.

| Protocol | Positive class | Negative class | cells x genes | # Positive | # Negative | Imbalance degree |
| --- | --- | --- | --- | --- | --- | --- |
| 10Xv2 | Natural killer cell | Cytotoxic T cell, CD4<sup>+</sup> T cell, CD14<sup>+</sup> monocyte, Megakaryocyte, B cell, CD16<sup>+</sup> monocyte, Dendritic cell, Plasmacytoid dendritic cell | (6444 x 22280) | 429 | 6015 | 0.929 |
| 10Xv2 | Cytotoxic T cell | Natural killer cell, CD4<sup>+</sup> T cell, CD14<sup>+</sup> monocyte, Megakaryocyte, B cell, CD16<sup>+</sup> monocyte, Dendritic cell, Plasmacytoid dendritic cell | (6444 x 22280) | 2128 | 4316 | 0.507 |
| 10Xv2 | CD4<sup>+</sup> T cell | Natural killer cell, Cytotoxic T cell, CD14<sup>+</sup> monocyte, Megakaryocyte, B cell, CD16<sup>+</sup> monocyte, Dendritic cell, Plasmacytoid dendritic cell | (6444 x 22280) | 1458 | 4986 | 0.708 |
| 10Xv3 | Natural killer cell | Cytotoxic T cell, CD4<sup>+</sup> T cell, CD14<sup>+</sup> monocyte, B cell, Megakaryocyte, CD16<sup>+</sup> monocyte, Dendritic cell | (3222 x 21905) | 194 | 3028 | 0.936 |
| 10Xv3 | Cytotoxic T cell | Natural killer cell, CD4<sup>+</sup> T cell, CD14<sup>+</sup> monocyte, B cell, Megakaryocyte, CD16<sup>+</sup> monocyte, Dendritic cell | (3222 x 21905) | 962 | 2260 | 0.574 |
| Drop-Seq | CD14<sup>+</sup> monocyte | Cytotoxic T cell, CD4<sup>+</sup> T cell, B cell, Natural killer cell, CD16<sup>+</sup> monocyte, Dendritic cell, Plasmacytoid dendritic cell, Megakaryocyte | (3222 x 19922) | 232 | 2990 | 0.922 |
| Drop-Seq | Cytotoxic T cell | CD14<sup>+</sup> monocyte, CD4<sup>+</sup> T cell, B cell, Natural killer cell, CD16<sup>+</sup> monocyte, Dendritic cell, Plasmacytoid dendritic cell, Megakaryocyte | (3222 x 19922) | 1189 | 2033 | 0.415 |
| Drop-Seq | CD4<sup>+</sup> T cell | CD14<sup>+</sup> monocyte, Cytotoxic T cell, B cell, Natural killer cell, CD16<sup>+</sup> monocyte, Dendritic cell, Plasmacytoid dendritic cell, Megakaryocyte | (3222 x 19922) | 889 | 2333 | 0.619 |
| inDrop | CD4<sup>+</sup> T cell | Cytotoxic T cell, CD14<sup>+</sup> monocyte, B cell, CD16<sup>+</sup> monocyte, Megakaryocyte, Plasmacytoid dendritic cell | (3222 x 17159) | 512 | 2710 | 0.811 |
| inDrop | Cytotoxic T cell | CD4<sup>+</sup> T cell, CD14<sup>+</sup> monocyte, B cell, CD16<sup>+</sup> monocyte, Megakaryocyte, Plasmacytoid dendritic cell | (3222 x 17159) | 1108 | 2114 | 0.476 |
| inDrop | CD14<sup>+</sup> monocyte | CD4<sup>+</sup> T cell, Cytotoxic T cell, B cell, CD16<sup>+</sup> monocyte, Megakaryocyte, Plasmacytoid dendritic cell | (3222 x 17159) | 1006 | 2216 | 0.546 |
| Seq-Well | CD4<sup>+</sup> T cell | Cytotoxic T cell, B cell, CD14<sup>+</sup> monocyte, Megakaryocyte, Dendritic cell, Plasmacytoid dendritic cell | (3176 x 21059) | 435 | 2741 | 0.841 |
| Seq-Well | Cytotoxic T cell | CD4<sup>+</sup> T cell, B cell, CD14<sup>+</sup> monocyte, Megakaryocyte, Dendritic cell, Plasmacytoid dendritic cell | (3176 x 21059) | 1121 | 2055 | 0.455 |

**Figure 4.**
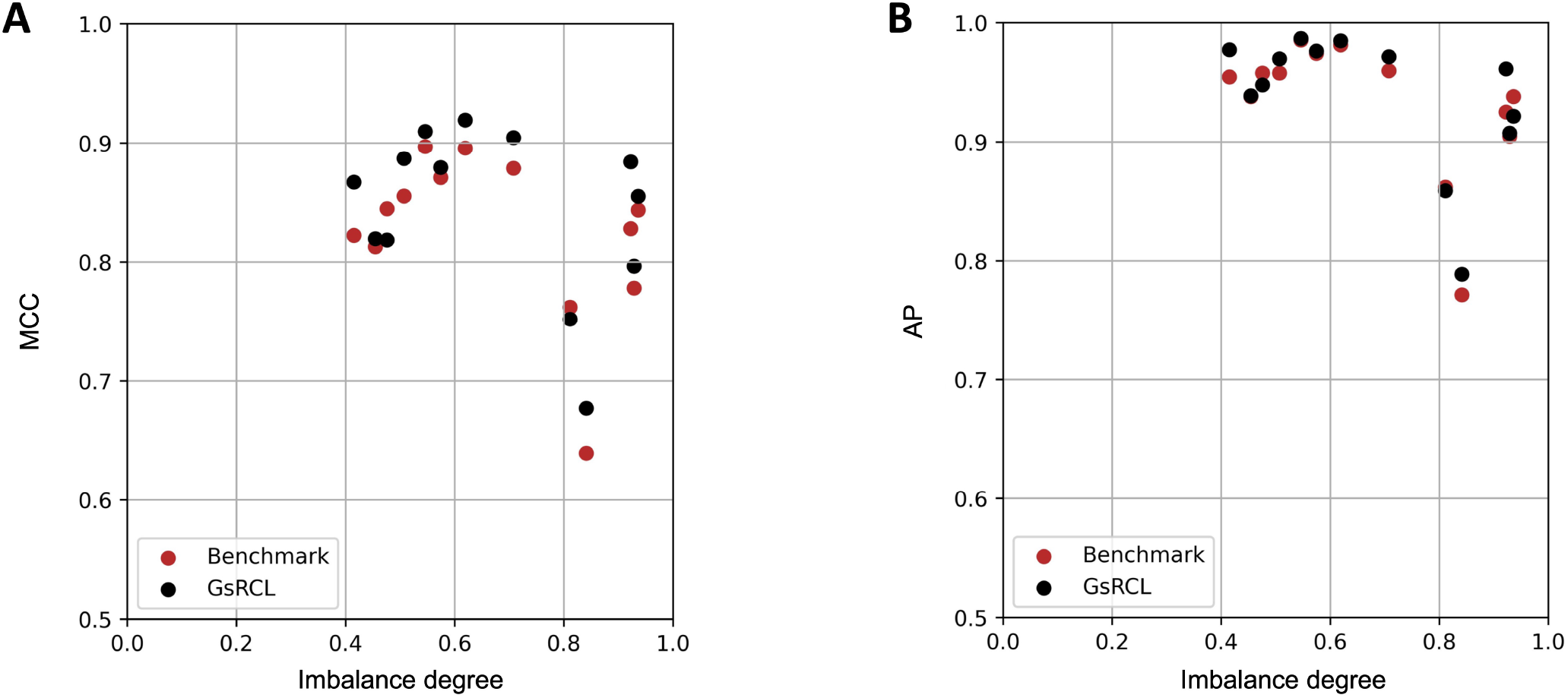
The relationships between the class imbalance degree and the predictive accuracy (i.e. MCC values (A) and AP values (B)) obtained by GsRCL and the benchmark method.

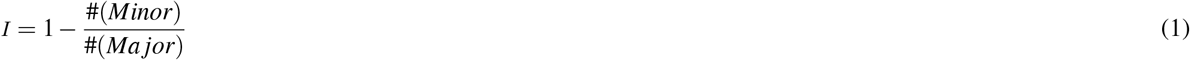

## Conclusion

In this work, we presented GsRCL – a novel Gaussian noise augmented contrastive learning-based method that learns a type of discriminative feature representations for the supervised cell-type identification tasks. The experimental results confirmed that GsRCL successfully improved the predictive accuracy of gene expression profiles-based cell-type identification method. Further analysis also revealed that self-supervised contrastive learning loss has stronger robustness against the overfitting issue w.r.t. down-stream supervised learning tasks. Future research could also focus on other data augmentation methods that would lead to better quality of positive and negative views in order to improve the performance of contrastive learning.

## Methods

### Self-supervised and supervised contrastive learning losses

In this work, we use both self-supervised and supervised contrastive learning losses to evaluate our proposed GsRCL method. Given a training batch with ℳ samples 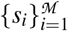, and let *Aug*^𝒩^(·) be a Gaussian noise-based augmentation method, for each instance *s*_*i*_, two views 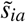 and 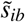 are generated to create a new batch with 2ℳ samples 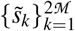, After transformations made by the encoder 𝒢 and the projector ℋ, i.e. *z* = ℋ (𝒢 (*Aug*^𝒩^(*s*))), the latent space representations of those two views of the target sample *s*_*i*_ are denoted by *z*_*a*_ and *z*_*b*_, which are also considered as a positive pair. The self-supervised contrastive learning loss^15^ is shown as Equation 2, where *sim*(*z*_*a*_, *z*_*b*_) is the cosine similarity and *τ* is a temperature parameter.

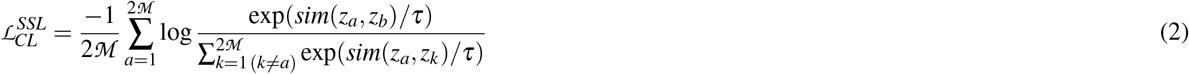

Analogously, the supervised contrastive learning loss also exploits the generated views as positive samples, but it further expand the positive set by considering the label information. For each augmented sample, the positive set will contain all samples that belong to the same class. All other samples that belong to a different class are considered negatives, as shown in Equation 3, where 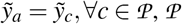, ∀*c* ∈ 𝒫, |𝒫| is the set of indices of all positives excluding *p* and |𝒫| is its cardinality.

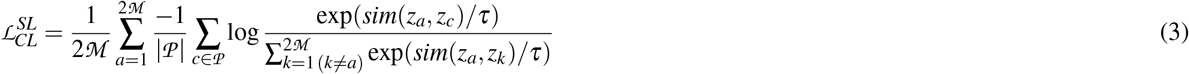

### Evaluating the predictive performance of the proposed GsRCL method

We evaluated the proposed GsRCL methods by using 16 single-cell RNA-seq datasets introduced in^8^. Due to the multi-class settings of those datasets, we transformed them into multiple binary classification problems using the well-known one-vs-the-rest strategy, i.e. each cell-type is treated as the positive class and all other cell-types form the negative class. Moreover, we further filtered out those datasets that consist of less than 30 testing instances for each fold of the 5-fold cross validation, and those datasets that lead to MCC values being greater than 0.90. Finally, 13 peripheral blood mononuclear cell (PBMC) datasets^33^ were used for the evaluation, as shown in Table 3. In addition, the gene expression profiles were further log-transformed. For the random genes masking data augmentation method, we followed the same approach in^29^. The evaluation is repeated using the top 500, 1000, 3000 and 5000 most variable genes. Consequently, our work involves the comparison between five augmentation methods, i.e. Gaussian noise-based augmentation, random masking 500, random masking 1000, random masking 3000, and random masking 5000.

We compared the predictive performance of the learned contrastive learning-based feature representations with the original scRNA-seq expression profiles by conducting a 5-fold cross validation for each dataset. Each of the original datasets were firstly split into two subsets, i.e. 80% of the instances were used to create a dataset for conducting the 5-fold cross validation, and the remaining 20% of the instances were used to create a validation-set for the contrastive learning networks training.

During the contrastive learning networks training stage, the validation-set was used to evaluate the predictive accuracy of the contrastive learning feature representations for every *n* epochs’ training. After every *n* epochs’ training, the contrastive learning networks were frozen and the encoding was used to generate contrastive learning feature representations for the original scRNA-seq expression profiles in the training folds and the validation-set. A SVM was used as the classifier to predict the class labels of the validation-set instances which were used to calculate the Matthews correlation coefficient (MCC) value. The optimal contrastive learning networks model was selected according to the highest MCC value, and then also used to generate contrastive learning feature representations for the test set. This training strategy is also adopted for using self-supervised and supervised contrastive losses working with the random genes masking data augmentation methods. We also used the original scRNA-seq expression profiles as a natural benchmark.

For the Gaussian noise-based augmentation method, we used a four-layer multi-layer perceptron as the encoder, three 4,096-dimensional hidden layers and a 2,048-dimensional output layer, i.e. representation layer. The projection head consists of a single 512-dimensional hidden layer and a 128-dimensional output layer. ReLU activation is used in both networks. The training is optimised using Adam with learning rate 10^−4^ and weight decay 10^−6^. The Gaussian noise used for augmentation is randomly drawn from a predefined Gaussian distribution 𝒩 (0, 1). For consistency, the same network architecture is used for the random genes masking data augmentation methods but with different sizes depending on the selected topmost variable genes. The batch size is set to 8,192 and the dropout ratio is set to 0.9. In terms of the training using two types of contrastive learning losses, we used 0.07 as the value of *τ*. The number of maximum training epochs is set to 3,000 and 1,000 for the self-supervised and supervised losses respectively. For the purpose of model selection, the validation performance is measured on every 40 epochs for the self-supervised loss and 20 epochs for the supervised loss. The grid-search was conducted for selecting the optimal hyperparamters of SVM during the second stage of GsRCL training – using the learned contrastive learning feature representations and a SVM to predict cell-types. The GsRCL method was implemented by PyTorch^34^ and Scikit-learn^35^.

### Availability and implementation

Source code is available via https://github.com/ibrahimsaggaf/GsRCL

## Acknowledgement

The authors acknowledge the academic research grants provided by Google Cloud. The authors also acknowledge the support by the Department of Computer Science and Information Systems and the Birkbeck GTA programme.

## References

1. Eberwine, J., Sul, J.-Y., Bartfai, T. & Kim, J. The promise of single-cell sequencing. Nat. methods 11, 25–27 (2014).

2. Edfors, F. et al. Gene-specific correlation of rna and protein levels in human cells and tissues. Mol. systems biology 12, 883 (2016).

3. de Boer, C. G. et al. Deciphering eukaryotic gene-regulatory logic with 100 million random promoters. Nat. Biotechnol. 38, 56–65 (2020).

4. Vaishnav, E. D. et al. The evolution, evolvability and engineering of gene regulatory dna. Nature 603, 455–463 (2022).

5. The modENCODE Consortium et al. Identification of functional elements and regulatory circuits by drosophila modencode. Science 330, 1787–1797 (2010).

6. Wan, C., Lees, J. G., Minneci, F., Orengo, C. & Jones, D. Analysis of temporal transcription expression profiles reveal links between protein function and developmental stages of drosophila melanogaster. PLOS Comput. Biol. 13, e1005791 (2017).

7. Plass, M. et al. Cell type atlas and lineage tree of a whole complex animal by single-cell transcriptomics. Science 360 (2018).

8. Abdelaal, T. et al. A comparison of automatic cell identification methods for single-cell rna sequencing data. Genome Biol. 20 (2019).

9. Hwang, B., Lee, J. H. & Bang, D. Single-cell rna sequencing technologies and bioinformatics pipelines. Exp. & molecular medicine 50, 1–14 (2018).

10. Plasschaert, L. W. et al. A single-cell atlas of the airway epithelium reveals the cftr-rich pulmonary ionocyte. Nature 560, 377–381 (2018).

11. Alquicira-Hernandez, J., Sathe, A., Ji, H. P., Nguyen, Q. & Powell, J. E. scpred: accurate supervised method for cell-type classification from single-cell rna-seq data. Genome biology 20, 1–17 (2019).

12. Nguyen, V. & Griss, J. scannotatr: framework to accurately classify cell types in single-cell rna-sequencing data. BMC bioinformatics 23, 1–13 (2022).

13. Ianevski, A., Giri, A. K. & Aittokallio, T. Fully-automated and ultra-fast cell-type identification using specific marker combinations from single-cell transcriptomic data. Nat. communications 13, 1–10 (2022).

14. Sun, Q., Peng, Y. & Liu, J. A reference-free approach for cell type classification with scrna-seq. Iscience 24, 102855 (2021).

15. Chen, T., Kornblith, S., Norouzi, M. & Hinton, G. A simple framework for contrastive learning of visual representations. In International conference on machine learning, 1597–1607 (PMLR, 2020).

16. He, K., Fan, H., Wu, Y., Xie, S. & Girshick, R. Momentum contrast for unsupervised visual representation learning. In IEEE/CVF Conference on Computer Vision and Pattern Recognition, 9729–9738 (2020).

17. Schneider, S., Baevski, A., Collobert, R. & Auli, M. wav2vec: Unsupervised pre-training for speech recognition. In Interspeech, 3465–3469 (2019).

18. Baevski, A., Schneider, S. & Auli, M. vq-wav2vec: Self-supervised learning of discrete speech representations. In The International Conference on Learning Representations, 1–12 (2019).

19. Khosla, P. et al. Supervised contrastive learning. In Larochelle, H., Ranzato, M., Hadsell, R., Balcan, M. & Lin, H. (eds.) Advances in Neural Information Processing Systems, vol. 33, 18661–18673 (2020).

20. Chen, T., Kornblith, S., Swersky, K., Norouzi, M. & Hinton, G. E. Big self-supervised models are strong semi-supervised learners. In Larochelle, H., Ranzato, M., Hadsell, R., Balcan, M. & Lin, H. (eds.) Advances in Neural Information Processing Systems, vol. 33, 22243–22255 (2020).

21. Kang, B., Li, Y., Xie, S., Yuan, Z. & Feng, J. Exploring balanced feature spaces for representation learning. In International Conference on Learning Representations (2021).

22. Robinson, J., Chuang, C.-Y., Sra, S. & Jegelka, S. Contrastive learning with hard negative samples. arXiv preprint 2010.04592 (2020).

23. Qu, Y. et al. Coda: Contrast-enhanced and diversity-promoting data augmentation for natural language understanding. arXiv preprint 2010.08670 (2020).

24. You, Y. et al. Graph contrastive learning with augmentations. In Larochelle, H., Ranzato, M., Hadsell, R., Balcan, M. & Lin, H. (eds.) Advances in Neural Information Processing Systems, vol. 33, 5812–5823 (2020).

25. Wang, Y., Wang, J., Cao, Z. & Barati Farimani, A. Molecular contrastive learning of representations via graph neural networks. Nat. Mach. Intell. 4, 279–287, DOI: https://doi.org/10.1038/s42256-022-00447-x (2022).

26. Xu, Y., Das, P. & McCord, R. P. Smile: mutual information learning for integration of single-cell omics data. Bioinformatics 38, 476–486 (2022).

27. Yan, X., Zheng, R. & Li, M. GLOBE: a contrastive learning-based framework for integrating single-cell transcriptome datasets. Briefings Bioinforma. 23 (2022). Bbac311.

28. Wang, X., Wang, J., Zhang, H., Huang, S. & Yin, Y. HDMC: a novel deep learning-based framework for removing batch effects in single-cell RNA-seq data. Bioinformatics 38, 1295–1303 (2021).

29. Ciortan, M. & Defrance, M. Contrastive self-supervised clustering of scrna-seq data. BMC bioinformatics 22, 1–27 (2021).

30. Wan, H., Chen, L. & Deng, M. scNAME: neighborhood contrastive clustering with ancillary mask estimation for scRNA-seq data. Bioinformatics 38, 1575–1583 (2022).

31. Han, W. et al. Self-supervised contrastive learning for integrative single cell RNA-seq data analysis. Briefings Bioinforma. 23 (2022). Bbac377.

32. Wan, C. & Freitas, A. A new hierarchical redundancy eliminated tree augmented naive bayes classifier for coping with gene ontology-based features. In Proceedings of the 33rd International Conference on Machine Learning (ICML 2016) Workshop on Computational Biology (2016).

33. Ding, J. et al. Systematic comparison of single-cell and single-nucleus rna-sequencing methods. Nat. biotechnology 38, 737–746, DOI: https://doi.org/10.1038/s41587-020-0465-8 (2020).

34. Paszke, A. et al. Pytorch: An imperative style, high-performance deep learning library. In Wallach, H. et al. (eds.) Advances in Neural Information Processing Systems 32, 8024–8035 (2019).

35. Pedregosa, F. et al. Scikit-learn: Machine learning in python. J. Mach. Learn. Res. 12, 2825–2830 (2011).

